# Comparison of Multi-objective Evolutionary Algorithms to Solve the Modular Cell Design Problem for Novel Biocatalysis

**DOI:** 10.1101/616078

**Authors:** Sergio Garcia, Cong Trinh

## Abstract

A large space of chemicals with broad industrial and consumer applications could be synthesized by engineered microbial biocatalysts. However, the current strain optimization process is prohibitively laborious and costly to produce one target chemical and often requires new engineering efforts to produce new molecules. To tackle this challenge, modular cell design based on a chassis strain that can be combined with different product synthesis pathway modules has been recently proposed. This approach seeks to minimize unexpected failure and avoid task repetition, leading to a more robust and faster strain engineering process. The modular cell design problem was mathematically formulated using a multi-objective optimization framework.[1] In this study, we evaluated a library of the state-of-the-art multi-objective evolutionary algorithms (MOEAs) to identify the most effective method to solve the modular cell design problem. Using the best MOEA, we found better solutions for modular cells compatible with many product synthesis modules. Furthermore, the best performing algorithm could provide better and more diverse design options that might help increase the likelihood of successful experimental implementation. We identified key parameter configurations to overcome the difficulty associated with multi-objective optimization problems with many competing design objectives. Interestingly, we found that MOEA performance with a real application problem, e.g., the modular strain design problem, does not always correlate with artificial benchmarks. Overall, MOEAs provide powerful tools to solve the modular cell design problem for novel biocatalysis.

## 1. Introduction

Multi-objective optimization is a powerful mathematical toolbox widely used in engineering disciplines to solve problems with multiple conflicting design objectives.[2] For example, in the field of chemical engineering, multi-objective optimization has been applied to balance design conflicts in the performance, material and energy requirements, and environmental sustainability of many different chemical processes.[3] In industrial biotechnology, with recent advancements in synthetic biology and metabolic engineering, microorganisms can be genetically modified to produce a large space of molecules with broad applications using renewable lignocellulosic biomass or waste products as feedstocks.[4,5] Due to the current strain design process that is prohibitively laborious and expensive [6], the application of modular design principles commonly used in engineering [7] to microbial biocatalysis has been recently proposed to overcome this challenge.[1,8–10] This modular cell design approach, known as ModCell, uses multi-objective optimization to account for the competing cellular objectives when cellular metabolism is (re)designed in a modular fashion to produce a diverse class of target chemicals. ModCell has been experimentally demonstrated for biosynthesis of alcohols [8,11,12] and esters [13–17] in *Escherichia coli*.

Despite the broad applicability of multi-objective optimization in engineering design, powerful solution algorithms remain elusive. Multi-objective evolutionary algorithms (MOEAs) are widely used techniques due to their flexibility and computational scalability.[18] MOEAs are based on a more general type of optimization method known as evolutionary algorithms, where candidate solutions, that represent individuals of a population, are iteratively modified using heuristic rules to increase their fitness. Recently, much attention has been placed in the development of MOEAs to solve many-objective problems (e.g., problems with 4 or more objectives) that often correspond to real-world applications, but can be very challenging to solve with conventional MOEAs.[19] For the case of ModCell problem, the popular MOEA NSGA-II[20,21] was used to design a modular cell under 20 different production modules,[1]. Due to the large chemical space of molecules that can potentially be synthesized by modular cells, scalability issues are expected to occur when constructing modular cells that are designed to be compatible with tenths or hundreds of products. Furthermore, using the best solver algorithm(s) allows to explore a more diverse design space, resulting in better choices for experimental implementation.

Many MOEAs have been proposed over the past two decades since the inception of landmark algorithms such as NSGAII [22] and SPEA2.[23] New MOEAs are benchmarked against libraries of artificial problems with known solutions, [24,25] and are expected to show enhanced performance for a subset of these problems in terms of scalability, identification of Pareto optimal solutions, and number of simulation generations needed to converge. This benchmarking methodology does not always reflect MOEA performance for general problems, since specialized parameter configurations or heuristics are often used and can lead to drastically different performance towards a specific problem of interest. Thus, the best MOEA for a certain application problem needs to be determined empirically. In this study, we evaluated a library of state-of-the-art MOEAs to solve the multi-objective ModCell problem, with special consideration for many-objective methods. Several cases study of increasing difficulty were examined using common performance indicators of solution optimality and diversity, and critical algorithm parameters that determine solution quality are also investigated.

## 2. Methods

### 2.1 Multi-objective modular cell design

Modular cell design enables rapid assembly of strains with desirable phenotypes from a modular (chassis) cell,[1] resembling the efficiency advantages of modular design in conventional engineering disciplines.[7,10] A modular cell is constructed by eliminating genes from a parent strain to maintain only core metabolic pathways shared across all pathway modules. Each module enables an optimized target product synthesis phenotype that leads to high yields, titers, and production rates. The different biochemical nature of each target metabolite can make the objectives compete with each other, turning the modular cell design problem into a multi-objective optimization problem known as ModCell2:[1]

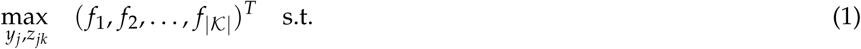

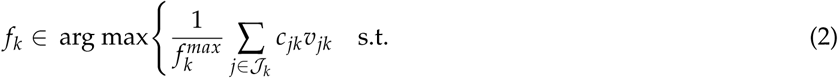

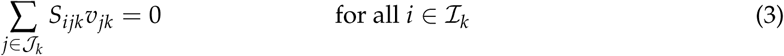

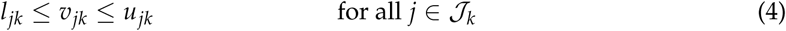

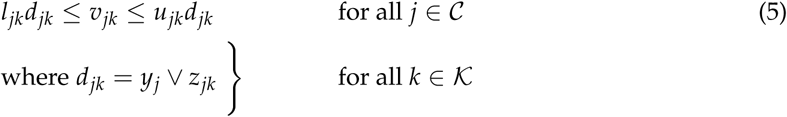

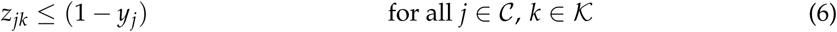

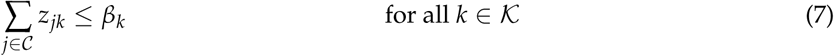

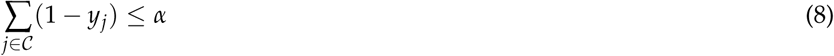

where 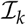, 𝒥_*k*_, and 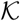 are the sets of metabolites, reactions, and associated production metabolic networks (i.e., the combination of the chassis organism with a specific product synthesis pathway), respectively. The desirable phenotype *f*_*k*_ for production module *k* (1) is determined based on key metabolic fluxes *v*_*jk*_ (mmol/gDCW/h) predicted by the constraint-based metabolic model [26] (2-5). For example, a common design objective is weak growth coupled to product formation (*wGCP*), that requires a high product synthesis rate at the maximum growth-rate, enabling growth selection of optimal production strains. Thus, in *wGCP* design, the inner optimization problem seeks to maximize growth rate through the linear objective function *c*_*jk*_ (2) subject to: (i) mass-balance constraints (3), where *S*_*ijk*_ represents the stoichiometric coefficient of metabolite *i* in reaction *j* of production network *k*, (ii) flux bound constraints (4) that determine reaction reversibility and available substrates, where *l*_*jk*_ and *u*_*jk*_ are lower and upper bounds respectively, and (iii) genetic manipulation constraints (5), i.e., deletion of a reaction *j* in the chassis through the binary indicator *y*_*j*_, or insertion of a reaction *j* in a specific production network *k* through the binary indicator *z*_*jk*_. The maximum product synthesis rate of each production network *k*, 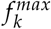, is determined by maximizing the product synthesis reaction subject to (3-4), allowing to bound *f*_*k*_ in *wGCP* between 0 and 1. Only a subset of all metabolic reactions, *C*, are considered as candidates for deletion, since many of the reactions in the metabolic model cannot be manipulated to enhance the target phenotype. Certain reactions can be deleted in the chassis but inserted back under specific production modules, enabling the chassis to be compatible with a broader number of modules (6). The number of module-reactions additions and reaction deletions in the chassis is constrained by parameters *β*_*k*_ (7) and *α* (8), respectively, to avoid unnecessary genetic manipulations that are generally time-consuming to implement and can lead to unforeseen phenotypes.

### 2.2 Optimal solutions for a multi-objective optimization problem

Optimal solutions for a multi-objective optimization problem (1-8) are defined based on the concept of domination: A vector *a* = (*a*_1_, …, *a*_*K*_)^*T*^ *dominates* another vector *b* = (*b*_1_, …, *b*_*K*_)^*T*^, denoted as *a* ≺ *b* if and only if *a*_*i*_ ≥ *b*_*i*_ ∀*i* ∈ {1, 2, …, *K*} and *a*_*i*_ = *b*_*i*_ for at least one *i*. Letting *x* be the design variables (i.e., *y*_*j*_ and *z*_*jk*_) and *X* be the feasible set determined by the problem constraints (2-8), a feasible solution *x*^∗^ ∈ *X* of the multi-objective optimization problem is called a Pareto optimal solution if and only if there does not exists a vector *x*′ ∈ *X* such that *F*(*x*′) ≺ *F*(*x*^*∗*^). The set of all Pareto optimal solutions is called Pareto set:

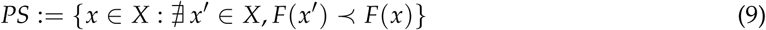

The projection of the Pareto set in the objective space is denoted as Pareto front:

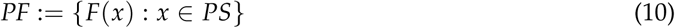

### 2.3 MOEA selection

To find the best MOEAs for ModCell2, we evaluated a recent and comprehensive set of MOEAs implemented in the PlatEMO platform.[27] From over 50 algorithms available in PlatEMO, we selected 2 methods for benchmark study, NSGAII/gamultiobj and MOEAIGDNS, and 8 methods that have been specifically developed to tackle many-objective problems with discrete variables like ModCell2, including ARMOEA, EFRRR, MaOEADDFC, SPEAR, tDEA, BiGE, NSGAIII, and SPEA2SDE (Table 1). It should be noted that gamultiobj is an alternative implementation of the NSGAII algorithm available in Matlab.

**Table 1.** Summary of MOEAs used in this study

| Abbreviation | Name | Notes | Reference |
| --- | --- | --- | --- |
| NSGAII | Non-dominated sorting genetic algorithm 2 | Highly applied MOEA | [22] |
| gamultiobj | Matlab implementation of NSGAII | Used in the original ModCell2 study[1] | [20] |
| MOEAIGDNS | Multi-objective evolutionary algorithm based on an enhanced inverted generational distance metric | General MOEA with an implementation that works well with discrete variables | [28] |
| ARMOEA | Adaptation to reference points multi-objective evolutionary algorithm | Many-objective EA based on MOEAIGDNS | [29] |
| EFRRR | Ensemble fitness ranking with ranking restriction | Many-objective EA | [30] |
| MaOEADDFC | Many-objective evolutionary algorithm based on directional diversity and favorable convergence | Many-objective EA | [31] |
| SPEAR | Strength Pareto evolutionary algorithm based on reference direction | Many-objective EA | [32] |
| tDEA | θ-dominance evolutionary algorithm | Many-objective EA | [33] |
| BiGE | Bi-goal evolution | Many-objective EA | [34] |
| NSGAIII | Non-dominated sorting genetic algorithm 3 | Many-objective EA | [35] |
| SPEA2SDE | Strength Pareto evolutionary algorithm 2 with shift-based density estimation | Many-objective EA | [36] |

### 2.4 Performance metrics

To evaluate the performance of different MOEAs for a given problem, each algorithm is ran for the same number of generations, and the resulting solutions, known as Pareto front approximations, are compared using functions that measure two qualities: (i) solution accuracy, i.e., to determine how similar the solution is to the true Pareto front and (ii) solution diversity, i.e., to evaluate how well distributed are the points in the solution. We selected the top 5 most used metrics according to a recent literature survey.[37] These include, in order of popularity, hypervolume (*HV*), generational distance (*GD*), epsilon indicator (*ϵ*), inverted generational distance (*IGD*), and coverage (*C*). Based on a recent study,[38] we considered the average Hausdorff distance (∆_*p*_), that combines *GD* and *IGD*, and hence simplified the number of performance metrics to 4 in our study. These metrics are defined as follows:

#### HV

This metric measures the volume occupied by the union of the smallest hyperboxes formed by each point in the Pareto front approximation and the reference point. This Pareto front approximation corresponds to the solution of a specific MOEA (denoted as *PF*) and the reference point is selected to be greater or equal to the maximum value attainable by any objective, which in our case is 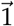 (Figure 1a):

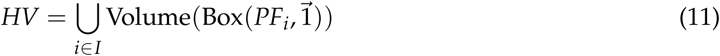

where *I* is the index set of *PF* points.

**Figure 1.**
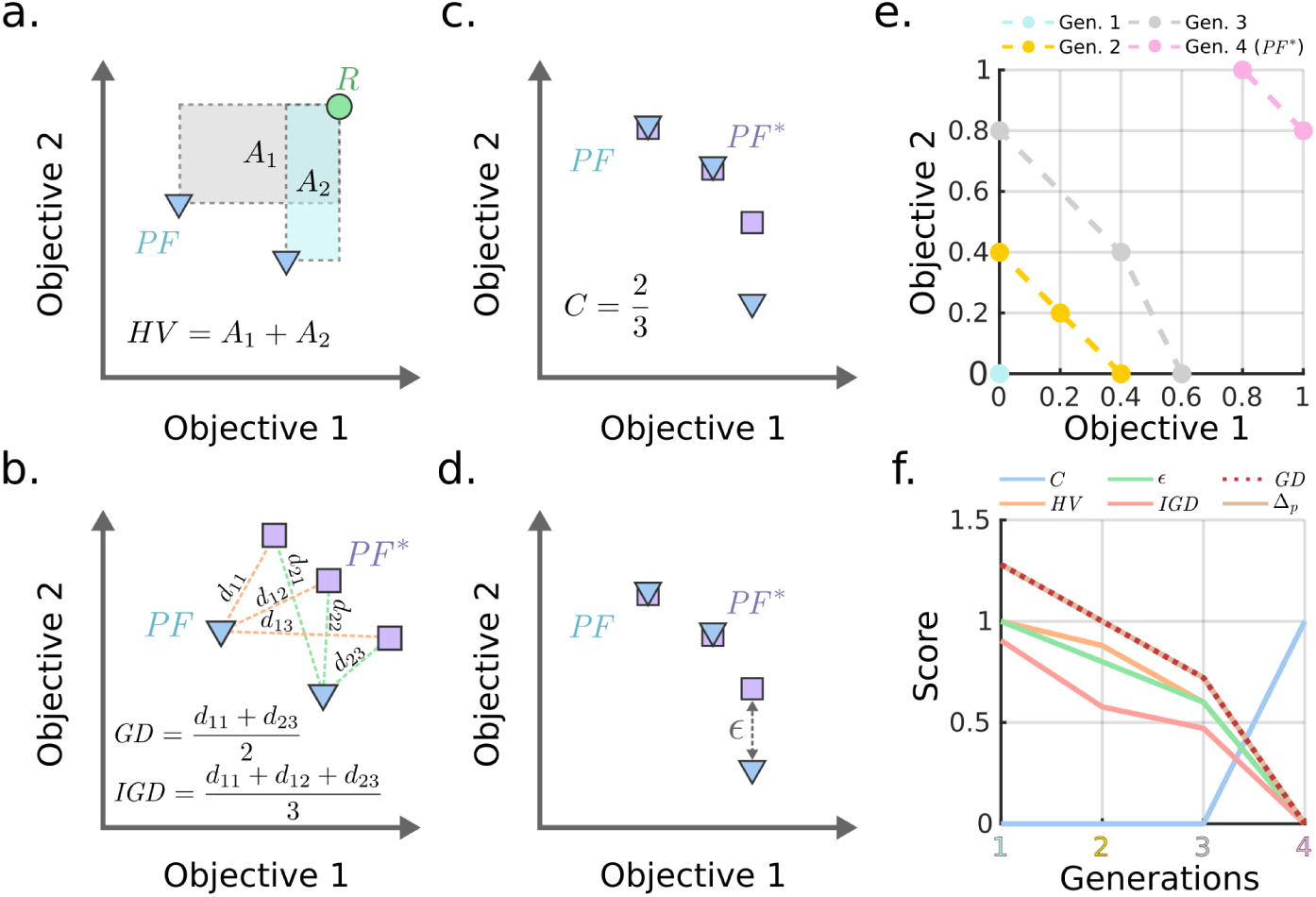
(**a-d**) Conceptual illustration of performance metrics of MOEAs for two-objective design problem. *PF* and *PF*^*∗*^ correspond to the Pareto front approximation and the best Pareto front available, respectively. The reference point *R* must always dominate all solutions in *PF*. (**e-f**) An example of Pareto fronts with 2 dimensions and associated metrics. The 4th generation corresponds to *PF*^*∗*^ used as a reference for comparison.

#### GD

This metric measures the distance between the solution *PF* and the best Pareto front approximation determined by combining non-dominated points from all MOEA solutions of a specific case study, denoted *PF*^*∗*^. More specifically, *GD* corresponds to the average Euclidean distance between each point in *PF* and the nearest point in *PF*^*∗*^, denoted as 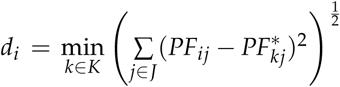, where *I*(*i* ∈ *I*), *K* (*k* ∈ *K*), and *J* (*j* ∈ *J*) correspond to the index sets of *PF* points, *PF*^*∗*^ points, and problem objectives, respectively (Figure 1b):

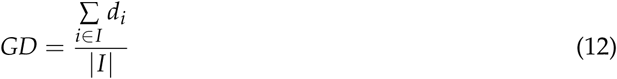

#### IGD

This metric measures the distance between *PF* and *PF*^*∗*^. It is determined by the average Euclidean distance between each point in *PF*^*∗*^ and the nearest point in *PF* denoted 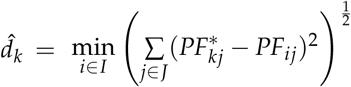 (Figure 1b):

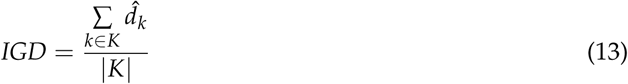

∆_*p*_: This metric combines *GD* and *IGD* metric and thus has superior properties:[38]

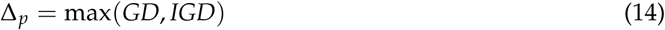

*C*: This metric determines the fraction of *PF*^*∗*^ captured by the solution *PF* (Figure 1c):

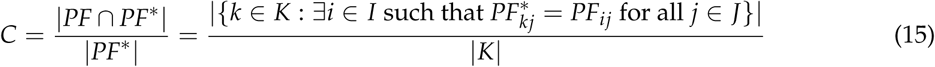

*ϵ*: This metric is the additive epsilon indicator[39] that measures the smallest value to be added to any point in *PF* to make it non-dominated with respect to some point in *PF*^*∗*^. In other words, it is the smallest value *ϵ* such that for any solution in *PF*^*∗*^ there is at least one solution in *PF* that is not worse by a difference of *ϵ* (Figure 1d):

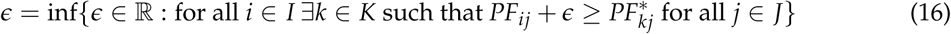

Use of these metrics can be illustrated with a two-objective design example with 4 generations of improving Pareto front approximations, where the final Pareto front is used as a reference (i.e., *PF*^*∗*^) (Figure 1e). As the Pareto fronts contain points that dominate the previous generations, all metrics decrease monotonically with the exception of *C* that increases to a value of 1 when both Pareto front approximation and reference are the same (Figure 1f).

### 2.5 Algorithm parameters

All parameters used in the simulations of this study were left as default except the following ones. The total number of generations was set to be 200, which is sufficient to reach high quality solutions for the problems of this study. In addition, the population size was set to be 100 for all algorithms unless noted otherwise. All problems were solved in triplicates with unique random number generator seeds.

### 2.6 Metabolic models

For all simulations, we used a core *E. coli* model, downloaded from the BiGG database[40] (https://bigg.ucsd.edu), that captures the most important metabolic pathways.[26] The product synthesis pathways for each module correspond to native *E. coli* pathways togheter with well-characterized heterologus pathways for the synthesis of propanol,[41] butanol,[42] isobutanol,[43] and pentanol.[41] The metabolic reactions associated with these pathways are described in the software implementation (Supplementary Material 1).

### 2.7 Implementation

The simulations were performed using the ModCell2 software framework.[1] The MOEAs are implemented in the PlatEMO Matlab library,[27] except *gamultiobj* which is implemented as part of the Matlab Optimization Toolbox. *HV* was calculated using the *hv package*.[44] All computations were executed in a computer with the Arch Linux operative system, Intel Core i7-3770 processor, and 32 GB of random-access memory. The Matlab 2018b code used to generate the results of this manuscript is available in Supplementary Material 1 and https://github.com/trinhlab/compare-moea.

## 3. Results and Discussion

### 3.1 Case 1: A 3-objectives design problem

We first formulated a design problem that considers an *E. coli* core model and 3 production modules based on the endogenous acetate, D-lactate, and ethanol biosynthesis pathways (Figure 2a). We used all MOEAs to solve for the problem by setting the *wGCP* design objective, a maximum number of reaction deletions *α* = 3, and no module reactions *β* = 0. These design parameters were sufficiently restrictive to generate conflicting objectives. A total coverage of *PF*^*∗*^ (*C* = 1) was reached within 20 generations by several algorithms (Figure 2b, e, h, i) and by *gamultiobj* after 150 generations (Figure 2k), while the remaining algorithms could not attain *C* values above 0.8 (Figure 2c, d, f, g, j, l). In particular, MaOEADDFC and BiGE obtained the worst *C*, *ϵ*, and ∆_*p*_ values (Figure 2m). Although *C*, *ϵ*, and ∆_*p*_ values of BiGE indicated inferior performance, this algorithm had the lowest *HV* since it generated only one point with a high objective value (Figure 2o). Due to the simplicity of the problem, every algorithm except MaOEADDFC, tDEA, and BiGE converged to very similar Pareto fronts (Figure 2n-x), and 5 of them reached *C* = 1, indicating convergence to the reference Pareto front (Figure 2y).

**Figure 2.**
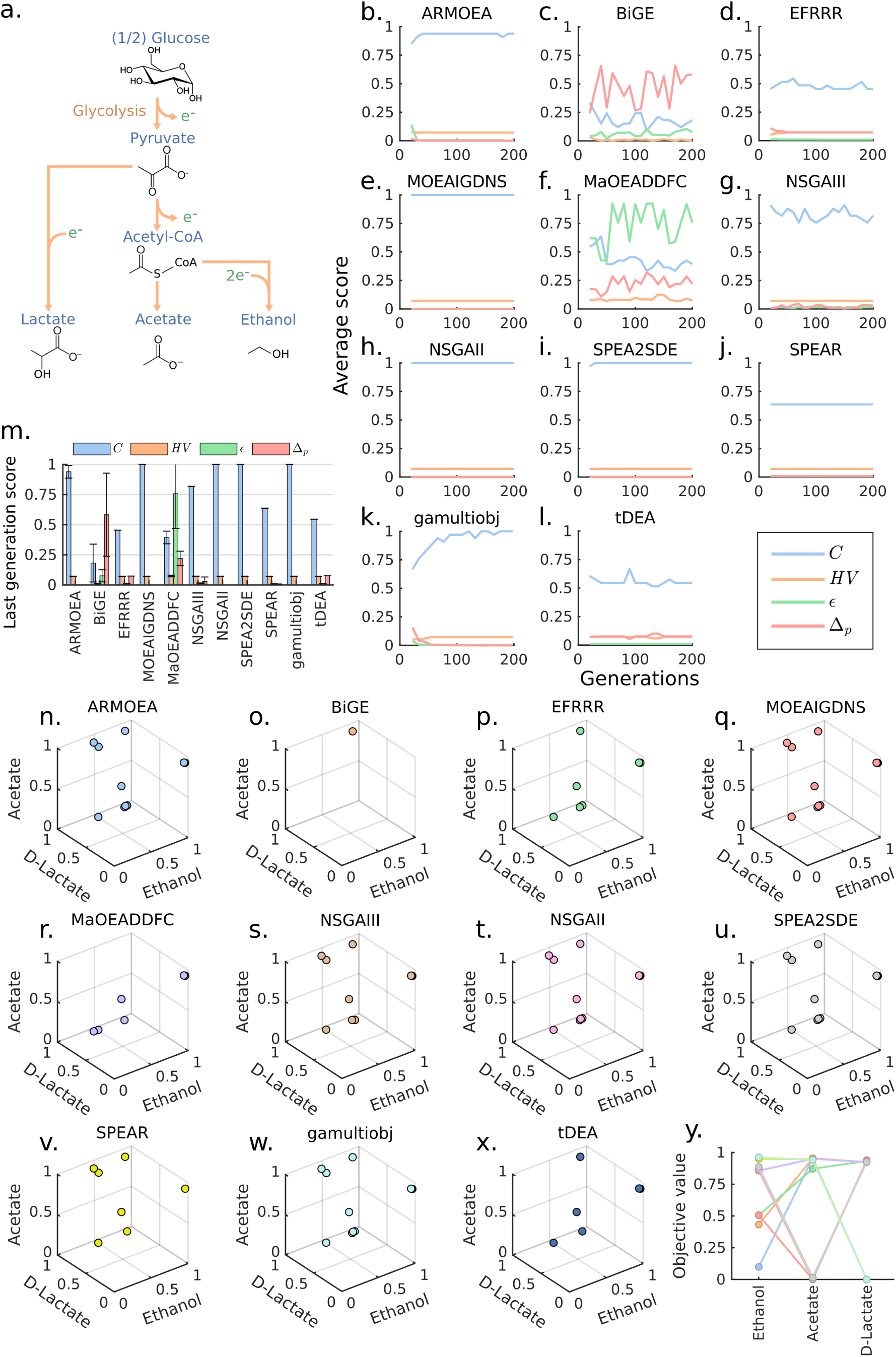
Comparison of MOEAs for a 3-objectives design problem. (**a**) The simplified metabolic pathways for conversion of glucose to the target products. Reducing equivalents are presented with *e*^*−*^. (**b-l**) Generation-dependent performance metrics for various MOEAs. (**m**) Performance metrics for various MOEAs at the last generation. (**n-x**) Pareto fronts of various MOEAs at the last generation. It should be noted that only the first replicate is plotted for clear illustration. (**y**) Reference Pareto front (*PF*^*∗*^). Each line represents a solution.

### 3.2 Case 2: A 10-objectives design problem

Using the same model and design parameters as in Case 1, we expanded the number of objectives to represent a more realistic scenario. These objectives correspond to 6 endogenous pathways for biosynthesis of D-lactate, acetate, ethanol, formate, pyruvate and L-glutamate and 4 heterologous pathways for biosynthesis of propanol, butanol, isobutanol, and pentanol. The additional objectives increased the difficulty of the problem, leading to more notable difference among algorithm performances (Figure 3a-k). The SPEA2SDE algorithm displayed consistent improvement of *C* as generations progressed, and quickly reached the smallest values of *ϵ* and ∆_*p*_ (Figure 3h). Other algorithms, including ARMOEA and MOEAIGDNS, also improved their *ϵ* with the increasing number of generations and reached the same final values of *ϵ* and ∆_*p*_ as SPEA2SDE (Figure 3a, d). However, SPEA2SDE approached *C* ≅ 0.6, which is twice the value reached by the next best-performing methods (Figure 3l). Remarkably, SPEA2SDE outperformed every other algorithm in all metrics, except *HV*. The *HV* metric continues to show bias towards algorithms that generated a small number of points and scored poorly in other metrics.

**Figure 3.**
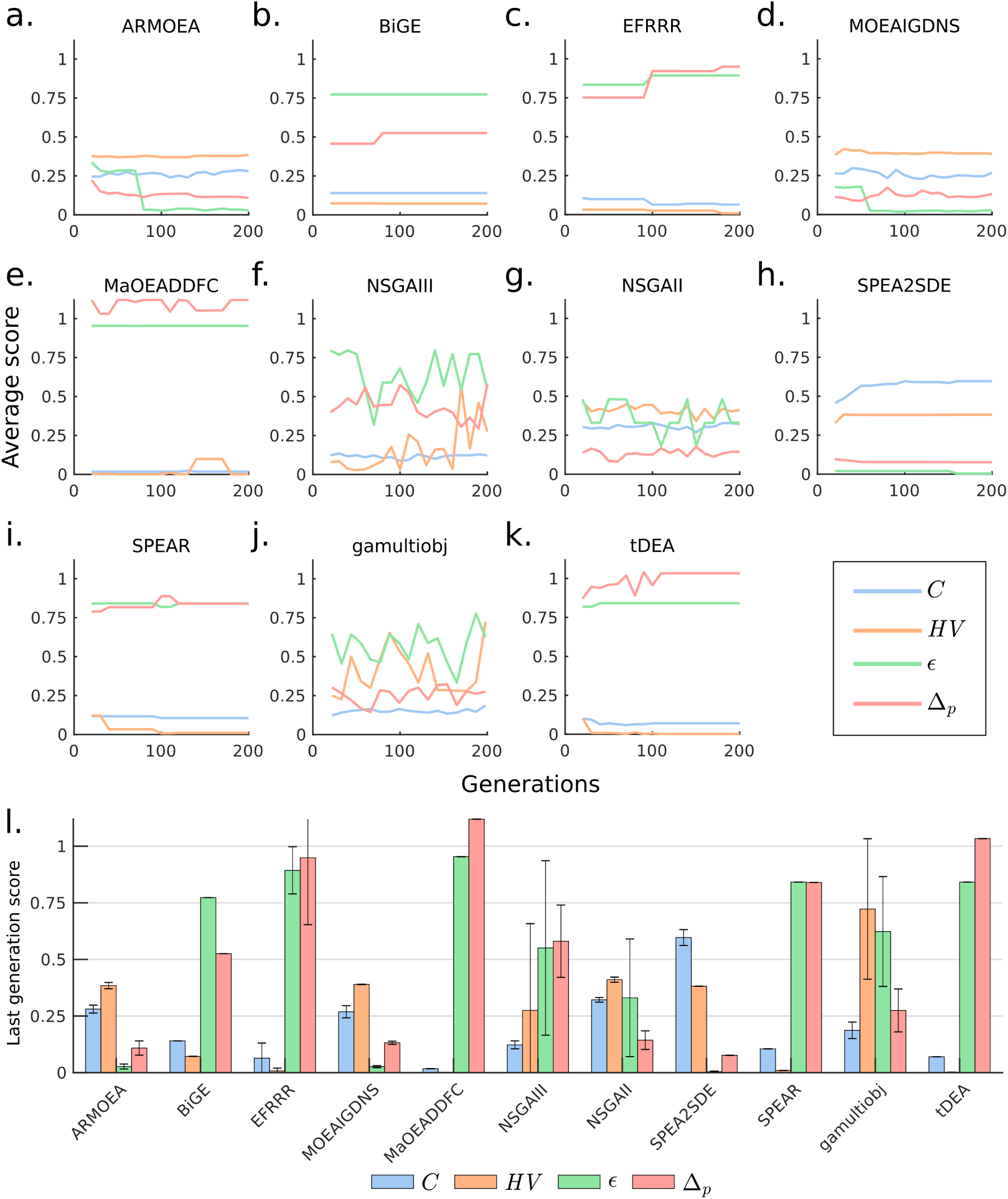
Comparison of MOEAs for a 10-objective design problem. (**a-k**) Generation-dependent performance metrics for various MOAEs. (**l**) Performance metrics for various MOEAs at the last generation.

### 3.3 Case 3: Use of large population size overcomes poor MOEA performance

Increasing the number of objectives often leads to a combinatorial explosion of the number of feasible Pareto optimal points and consequently causes poor MOEA performance. This problem can be alleviated by using a larger population size to sample a broader volume of solution space.[45] To test this strategy for the 10-objectives design problem above, we increased the population size from 100 to 1000 individuals. The result showed that ARMOEA, MOEAIGDNS, NSGAII, SPEA2SDE (the best performer in Case 2), and *gamultiobj*, could reach *C* of 0.7, *ϵ* of 0, and ∆_*p*_ of 0 in fewer than 50 generations (Figure 4a, d, g, h, j). These 5 algorithms also yielded very similar final values across all metrics (Figure 4l). The remaining algorithms converged to considerably lower *C* values (Figure 4b, c, e, f, i, k). Remarkably, NSGAII/*gamultiobj*, that is not considered a many-objective solver, performed better than more recent many-objective algorithms such as NSGAIII.

**Figure 4.**
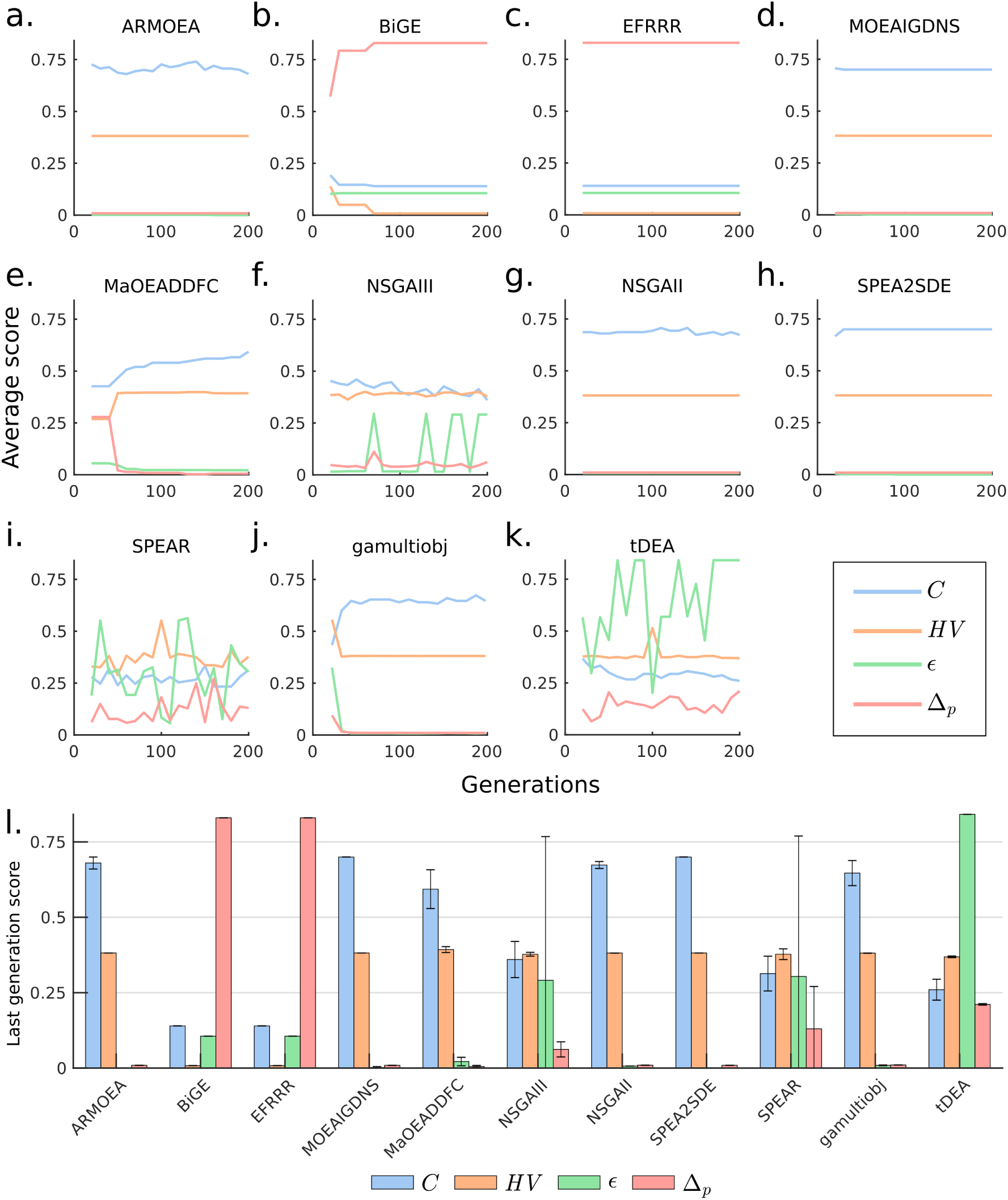
Comparison of MOEAs for a 10-objective design problem with larger population sizes (**a-k**) Generation-dependent performance metrics for various MOAEs. (**l**) Performance metrics for various MOEAs at the last generation.

One limitation of using larger populations is an increased cost in computational time. We observed that a 10-fold increase in population sizes resulted in a 10-fold increase in the run times (Figure 5). Nonetheless, all metrics reached a stable value in the top performing algorithms after 50 generations (out of 200 total), suggesting that fewer generations were needed by using a larger population size. Among the best performing algorithms with large population sizes, *gamultiobj*, implemented in the Matlab Optimization Toolbox, required the shortest run time, followed by NSGAII and SPEA2SDE implemented in PlatEMO.

**Figure 5.**
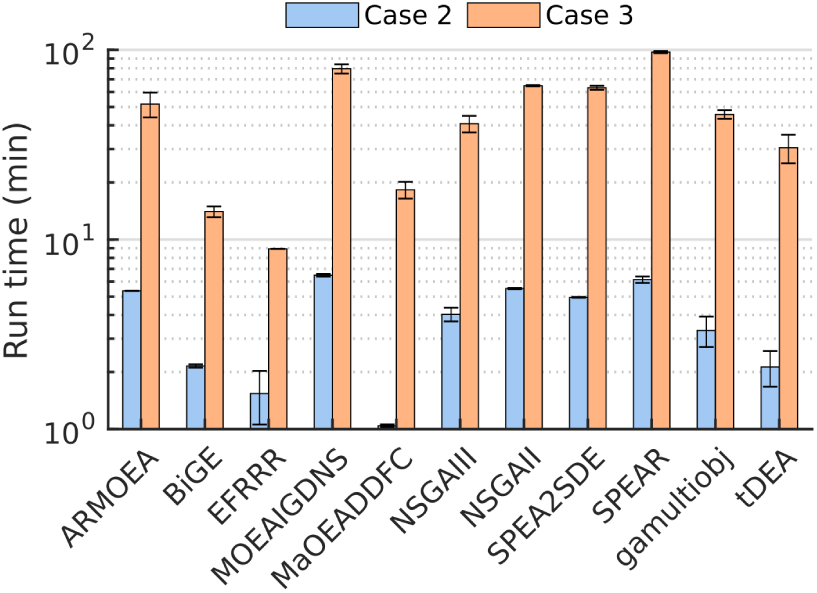
Wall-clock run times for the 10-objective design problem with population sizes of 100 (Case 2) and 1000 (Case 3).

## 4. Conclusions

In this study, we evaluated the performance of several MOEAs to solve the modular cell design problem. SPEA2SDE, the recently developed many-objective method, was the best performing MOEA under limited population sizes in our study. However, for sufficiently large populations, several algorithms attained the best results, including the well-established NSGAII, which performed better under large populations than more recently developed many-objective MOEAs. We used the most popular performance metrics to compare MOEAs and found that the coverage (*C*) metric is the most valuable indicator. This metric can provide an intuitive quantitative meaning and tends to increase monotonically with the number of generations simulated. In contrast, hypervolume (*HV*) generally did not differentiate algorithm performance and was misleading in some scenarios where an algorithm generated very few solutions. Overall, these results highlight the need for empirical testing of MOEAs towards specific problems and of the population size as a more important factor in performance than the unique heuristics used by different algorithms.

## Author Contributions

CTT initiated and supervised the study. SG and CTT designed experiments. SG performed simulation experiments and analyzed data. SG and CTT wrote and approved the manuscript.

## References

1. Garcia, S.; Trinh, C. Multiobjective strain design: A framework for modular cell engineering. Metabolic Engineering 2019, 51. doi:10.1016/j.ymben.2018.09.003.

2. Coello, C.A.C.; Lamont, G.B. Applications of multi-objective evolutionary algorithms; Vol. 1, World Scientific, 2004.

3. Rangaiah, G.P. Multi-objective optimization: techniques and applications in chemical engineering; Vol. 1, World Scientific, 2009.

4. Trinh, C.T.; Mendoza, B. Modular cell design for rapid, efficient strain engineering toward industrialization of biology. Current Opinion in Chemical Engineering 2016, 14, 18–25.

5. Lee, S.Y.; Kim, H.U.; Chae, T.U.; Cho, J.S.; Kim, J.W.; Shin, J.H.; Kim, D.I.; Ko, Y.S.; Jang, W.D.; Jang, Y.S. A comprehensive metabolic map for production of bio-based chemicals. Nature Catalysis 2019, 2, 18.

6. Nielsen, J.; Keasling, J. Engineering Cellular Metabolism. Cell 2016, 164, 1185–1197. doi:10.1016/j.cell.2016.02.004.

7. Bonvoisin, J.; Halstenberg, F.; Buchert, T.; Stark, R. A systematic literature review on modular product design. Journal of Engineering Design 2016, 27, 488–514. doi:10.1080/09544828.2016.1166482.

8. Trinh, C.T. Elucidating and reprogramming Escherichia coli metabolisms for obligate anaerobic n-butanol and isobutanol production. Applied microbiology and biotechnology 2012, 95, 1083–1094.

9. Trinh, C.T.; Liu, Y.; Conner, D.J. Rational design of efficient modular cells. Metabolic engineering 2015, 32, 220–231.

10. Garcia, S.; Trinh, C. Modular design: Applying proven engineering principles to biotechnology. Under review 2019.

11. Trinh, C.T.; Li, J.; Blanch, H.W.; Clark, D.S. Redesigning Escherichia coli metabolism for anaerobic production of isobutanol. Appl. Environ. Microbiol. 2011, 77, 4894–4904.

12. Wilbanks, B.; Layton, D.; Garcia, S.; Trinh, C. A Prototype for Modular Cell Engineering. ACS Synthetic Biology 2017, p. acssynbio.7b00269. doi:10.1021/acssynbio.7b00269.

13. Layton, D.S.; Trinh, C.T. Engineering modular ester fermentative pathways in Escherichia coli. Metabolic Engineering 2014, 26, 77–88. doi:10.1016/j.ymben.2014.09.006.

14. Layton, D.S.; Trinh, C.T. Expanding the modular ester fermentative pathways for combinatorial biosynthesis of esters from volatile organic acids. Biotechnology and bioengineering 2016.

15. Layton, D.S.; Trinh, C.T. Microbial synthesis of a branched-chain ester platform from organic waste carboxylates. Metabolic Engineering Communications 2016, 3, 245–251.

16. Wierzbicki, M.; Niraula, N.; Yarrabothula, A.; Layton, D.S.; Trinh, C.T. Engineering an Escherichia coli platform to synthesize designer biodiesels. Journal of biotechnology 2016, 224, 27–34.

17. Lee, J.; Trinh, C.T. De novo Microbial Biosynthesis of a Lactate Ester Platform. bioRxiv 2018, p. 498576.

18. Marler, R.T.; Arora, J.S. Survey of multi-objective optimization methods for engineering. Structural and multidisciplinary optimization 2004, 26, 369–395.

19. Li, B.; Li, J.; Tang, K.; Yao, X. Many-objective evolutionary algorithms: A survey. ACM Computing Surveys (CSUR) 2015, 48, 13.

20. Matlab documentation gamultiobj Algorithm. https://www.mathworks.com/help/gads/gamultiobj-algorithm.html. Accessed: 2019-02-04.

21. Kalyanmoy, D. Multi objective optimization using evolutionary algorithms; John Wiley and Sons, 2001. Chichester, England.

22. Deb, K.; Pratap, A.; Agarwal, S.; Meyarivan, T. A fast and elitist multiobjective genetic algorithm: NSGA-II. IEEE transactions on evolutionary computation 2002, 6, 182–197.

23. Zitzler, E.; Laumanns, M.; Thiele, L. SPEA2: Improving the strength Pareto evolutionary algorithm. TIK-report 2001, 103.

24. Zitzler, E.; Deb, K.; Thiele, L. Comparison of multiobjective evolutionary algorithms: Empirical results. Evolutionary computation 2000, 8, 173–195.

25. Deb, K.; Thiele, L.; Laumanns, M.; Zitzler, E. Scalable multi-objective optimization test problems. Proceedings of the 2002 Congress on Evolutionary Computation. IEEE, 2002, Vol. 1, pp. 825–830.

26. Palsson, B.Ø. Systems biology: constraint-based reconstruction and analysis; Cambridge University Press, 2015.

27. Tian, Y.; Cheng, R.; Zhang, X.; Jin, Y. PlatEMO: A MATLAB platform for evolutionary multi-objective optimization. IEEE Computational Intelligence Magazine 2017, 12, 73–87.

28. Tian, Y.; Zhang, X.; Cheng, R.; Jin, Y. A multi-objective evolutionary algorithm based on an enhanced inverted generational distance metric. IEEE Congress on Evolutionary Computation (CEC). IEEE, 2016, pp. 5222–5229.

29. Tian, Y.; Cheng, R.; Zhang, X.; Cheng, F.; Jin, Y. An indicator-based multiobjective evolutionary algorithm with reference point adaptation for better versatility. IEEE Transactions on Evolutionary Computation 2018, 22, 609–622.

30. Yuan, Y.; Xu, H.; Wang, B.; Zhang, B.; Yao, X. Balancing convergence and diversity in decomposition-based many-objective optimizers. IEEE Transactions on Evolutionary Computation 2016, 20, 180–198.

31. Cheng, J.; Yen, G.G.; Zhang, G. A many-objective evolutionary algorithm with enhanced mating and environmental selections. IEEE Transactions on Evolutionary Computation 2015, 19, 592–605.

32. Jiang, S.; Yang, S. A strength Pareto evolutionary algorithm based on reference direction for multiobjective and many-objective optimization. IEEE Transactions on Evolutionary Computation 2017, 21, 329–346.

33. Yuan, Y.; Xu, H.; Wang, B.; Yao, X. A new dominance relation-based evolutionary algorithm for many-objective optimization. IEEE Transactions on Evolutionary Computation 2016, 20, 16–37.

34. Li, M.; Yang, S.; Liu, X. Bi-goal evolution for many-objective optimization problems. Artificial Intelligence 2015, 228, 45–65.

35. Deb, K.; Jain, H. An evolutionary many-objective optimization algorithm using reference-point-based nondominated sorting approach, part I: Solving problems with box constraints. IEEE Transactions on Evolutionary Computation 2014, 18, 577–601.

36. Li, M.; Yang, S.; Liu, X. Shift-based density estimation for Pareto-based algorithms in many-objective optimization. IEEE Transactions on Evolutionary Computation 2014, 18, 348–365.

37. Riquelme, N.; Von Lücken, C.; Baran, B. Performance metrics in multi-objective optimization. Latin American Computing Conference (CLEI). IEEE, 2015, pp. 1–11.

38. Schutze, O.; Esquivel, X.; Lara, A.; Coello, C.A.C. Using the averaged Hausdorff distance as a performance measure in evolutionary multiobjective optimization. IEEE Transactions on Evolutionary Computation 2012, 16, 504–522.

39. Zitzler, E.; Thiele, L.; Laumanns, M.; Fonseca, C.M.; Da Fonseca Grunert, V. Performance assessment of multiobjective optimizers: An analysis and review. TIK-Report 2002, 139.

40. King, Z.A.; Lu, J.; Dräger, A.; Miller, P.; Federowicz, S.; Lerman, J.A.; Ebrahim, A.; Palsson, B.O.; Lewis, N.E. BiGG Models: A platform for integrating, standardizing and sharing genome-scale models. Nucleic acids research 2015, 44, D515–D522.

41. Tseng, H.C.; Prather, K.L. Controlled biosynthesis of odd-chain fuels and chemicals via engineered modular metabolic pathways. Proceedings of the National Academy of Sciences 2012, p. 201209002.

42. Shen, C.R.; Lan, E.I.; Dekishima, Y.; Baez, A.; Cho, K.M.; Liao, J.C. High titer anaerobic 1-butanol synthesis in Escherichia coli enabled by driving forces. Applied and environmental microbiology 2011.

43. Atsumi, S.; Hanai, T.; Liao, J.C. Non-fermentative pathways for synthesis of branched-chain higher alcohols as biofuels. nature 2008, 451, 86.

44. Fonseca, C.M.; Paquete, L.; López-Ibánez, M. An improved dimension-sweep algorithm for the hypervolume indicator. IEEE international conference on evolutionary computation. IEEE, 2006, pp. 1157–1163.

45. Ishibuchi, H.; Sakane, Y.; Tsukamoto, N.; Nojima, Y. Evolutionary many-objective optimization by NSGA-II and MOEA/D with large populations. IEEE International Conference on Systems, Man and Cybernetics. IEEE, 2009, pp. 1758–1763.

